# COTAN: Co-expression Table Analysis for scRNA-seq data

**DOI:** 10.1101/2020.05.11.088062

**Authors:** S. G. Galfrè, F. Morandin, M. Pietrosanto, F. Cremisi, M. Helmer-Citterich

## Abstract

Estimating co-expression of cell identity factors in single-cell transcriptomes is crucial to decode new mechanisms of cell state transition. Due to the intrinsic low efficiency of single-cell mRNA profiling, novel computational approaches are required to accurately infer gene co-expression in a cell population. We introduce COTAN, a statistical and computational method to analyze the co-expression of gene pairs at single cell level, providing the foundation for single-cell gene interactome analysis.

## 1 Introduction

Single-cell transcriptomes can describe known cell identity states and uncover new ones. This is basically achieved by methods of analysis of correlation of RNA counts among cells and subsequent clustering of cells with consistent global gene expression^1, 2^. The typical pipeline requires to normalize and log-transform raw read counts, yielding expression levels, and perform multivariate analysis^3, 4^ on the latter. Unfortunately, the intrinsic low efficiency of scRNA-seq^5–7^ precludes the detection of many expressed genes in single cells, in particular in droplet-based experiments. This has a critical effect on expression levels, causing the appearance of dropouts artefacts^8, 9^, and often limiting the scope of the research to few tools based on zero-inflation and imputation^10–12^. However, the introduction of Unique Molecular Identifiers^13^ (UMI) greatly reduces amplification noise, and the resulting UMI counts typically fit simple probabilistic models, thus allowing approaches not based on normalization^8^. To exploit this feature with a focus on genes joint distribution, we introduce CO-expression Tables ANalysis (COTAN), which uses UMI count matrices without normalization and does not depend on zero-inflation.

COTAN is based on three main elements: a robust estimation of the UMI detection efficiency (UDE) of each cell, a flexible model for the probability of zero UMI counts, and a generalized contingency table framework for zero/non-zero UMI counts for couples of genes. In fact, COTAN estimates the co-expression of gene pairs by comparing the number of cells that have zero UMI counts for both genes, with the expected value under independence hypothesis. This approach is based on the fact that technical zeros are always independently distributed, while biological zeros may be correlated for genes associated to cell differentiation, providing a way to recover information on the joint distribution.

To estimate the model parameters, COTAN uses raw UMI counts, but then, for computing co-expressions these are coded as zero/non-zero. This choice is a key feature of COTAN and has the twofold aim to increase the sensitivity of the method for genes with low expression level and to better handle genes whose counts deviate from the chosen parametrical distributions.

COTAN can effectively assess co-expression of gene pairs, yielding for each pair a numerical index and an approximate *p*-value; it can investigate whether single genes are differentially expressed in the population, by scoring them with a global index of differentiation; finally, it provides ways to plot and cluster genes according to their co-expression pattern with other genes, effectively helping the study of gene interactions.

Though the paper is mostly self-contained, with a significant section of methods, the mathematical models are only drafted here: proofs and finer details can be found in the twin paper^14^.

## 2 Results

### 2.1 Main features

COTAN analysis is performed on post-quality-control UMI counts. It starts with the estimation of the model parameters: UDE for cells (denoted by *ν*_*c*_ in the Methods Section), and mean (*λ*_*g*_) and dispersion (*a*_*g*_) for the expression of genes. While UDEs and means are estimated by moment methods, for dispersions we fit the number of cells showing zero UMI counts with its expected value under the negative binomial model. This is done to improve the method robustness, as afterwards we only use dispersions to compute the probability of zero UMI counts. These probabilities allow to devise an independence test for the expression of two genes (gene-pair analysis, GPA). The test is based on generalized 2 × 2 contingency tables (also indicated as co-expression tables) which collect the joint occurrence of zero UMI counts for two genes.

COTAN provides an approximate *p*-value for the GPA test, and a signed co-expression index (COEX), which measures the direction and significance of the deviation from the independence hypothesis. Positive values denote joint expression of two genes, negative values denote disjoint expression and values near 0 are expected when one or both are constitutive genes, or when the statistical power is too low (Figure 1A, 2B).

**Figure 1.**
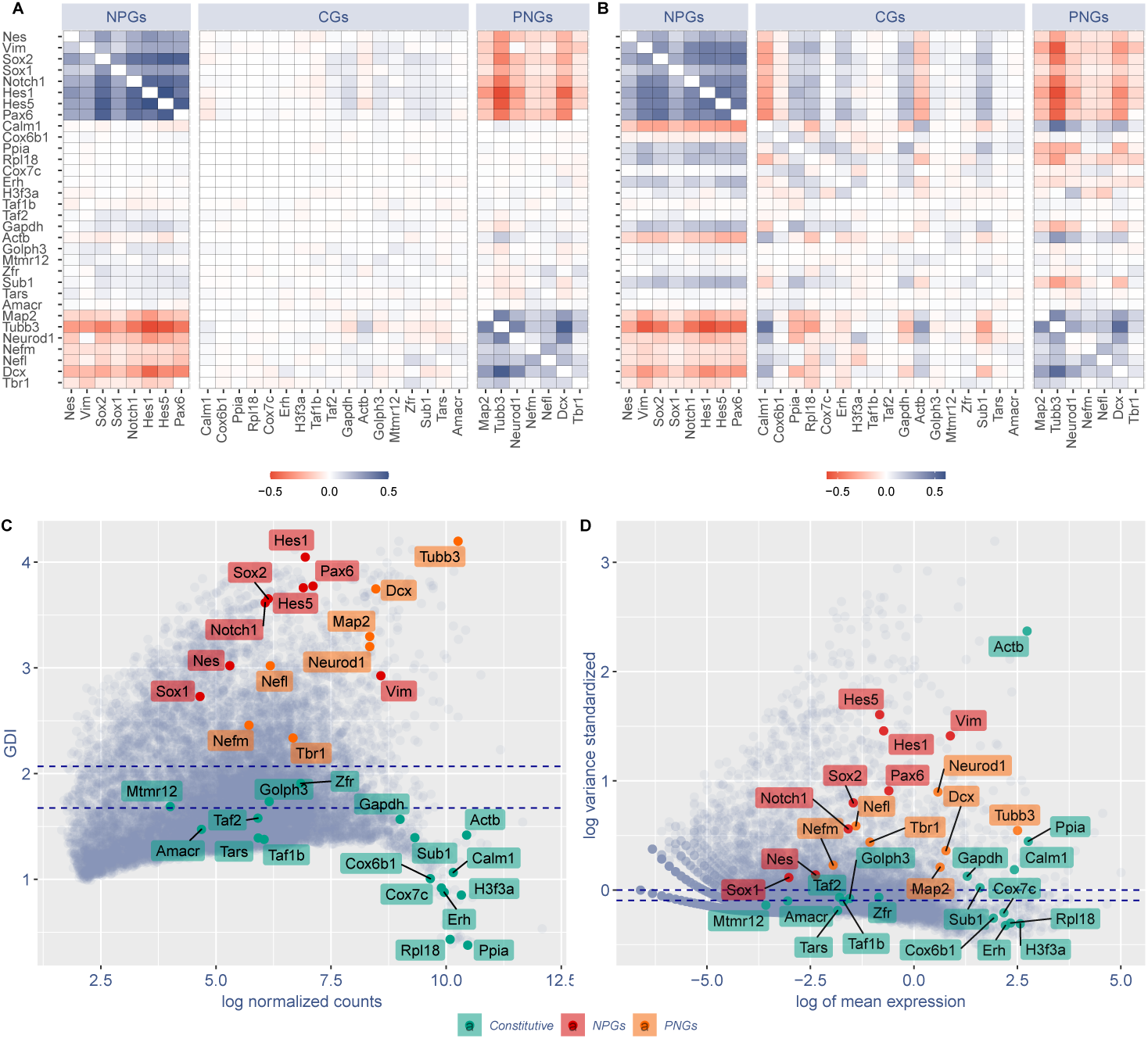
GPA and GDI are able to discriminate constitutive genes (CGs) from neural progenitor genes (NPGs) and pan-neuronal genes (PNGs). **A**. COTAN GPA of selected genes. Cell color encodes COEX index: blue indicates genes showing joint expression, red indicates genes showing disjoint expression. White indicates independence, meaning that one or both genes are constitutive, or that the statistical power is too low to detect co-expression (See Methods). Since the co-expression of a gene with itself is irrelevant, the diagonal is made artificially white. **B**. Pearson correlation matrix of the same selected genes as in (A), using Seurat^15^ normalized expression levels (obtained following the website vignettes – *Guided Clustering Tutorial*). **C, D**. Comparison between COTAN global differentiation index (GDI, C) and Seurat *highly variable features* (D) analysis. Red labels indicate NPGs, orange labels PNGs, green labels CGs. Dotted blue lines correspond to the median (lowest) and the third quartile (highest). All plots refer to E16.5 mouse hippocampal cells^16^ and genes are selected to be characteristic of NPGs, PNGs and constitutive genes with both high and low typical expression.

**Figure 2.**
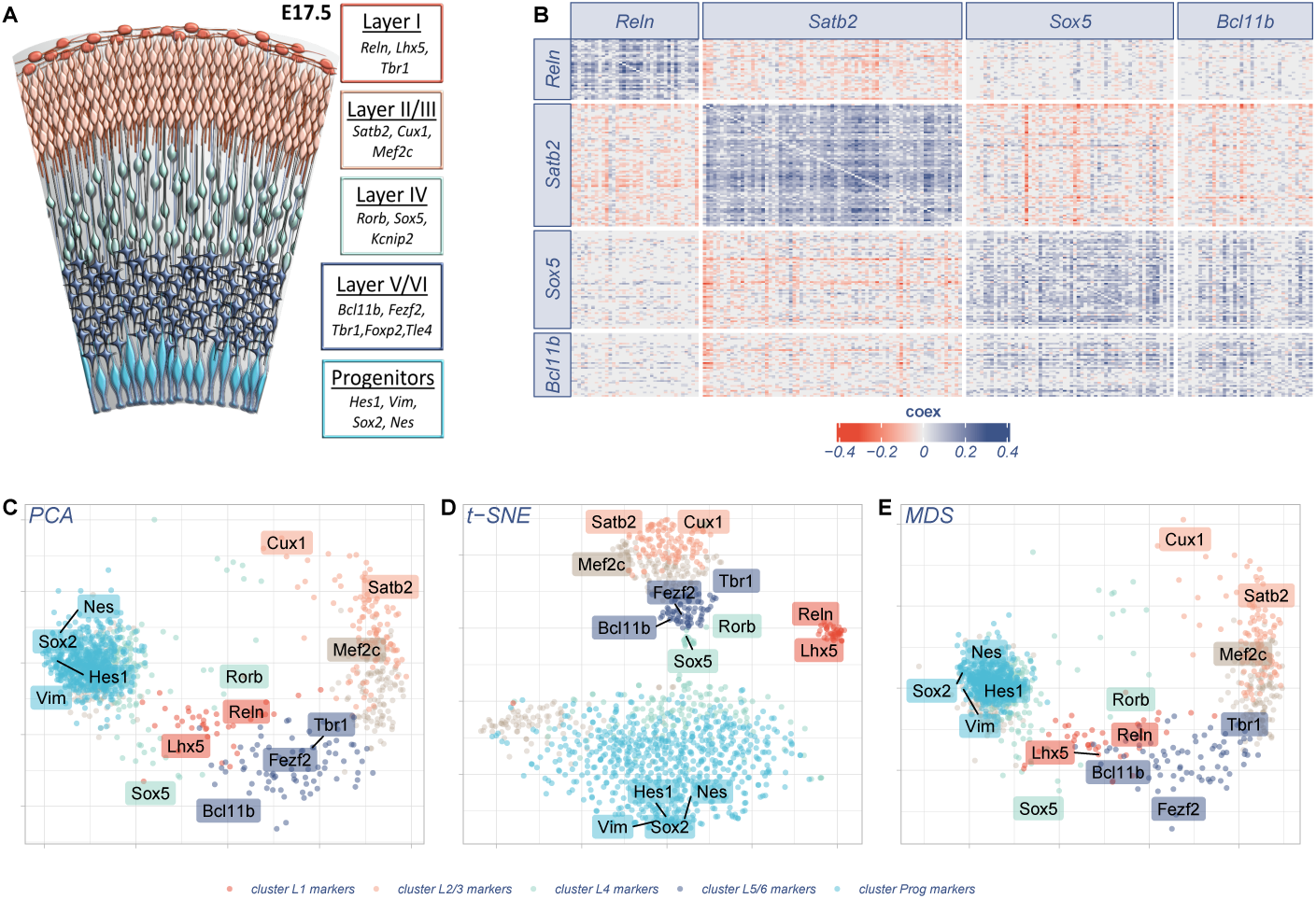
Gene clustering in scRNA-seq analysis. **A**. The six layers of differentiated neurons and progenitor cells of late embryonic cortex are depicted in different colors together with known markers of cell identity^18, 21^. **B**. GPA heatmap of the 248 genes showing strong joint expression with the genes indicated in labels: *Reln, Satb2, Sox5* and *Bcl11b* respectively markers of layers I, II/III, IV and V/VI. The heatmap shows the reciprocal relationship between all genes pairs; significant joint expression is indicated with blue (positive COEX values) while significant disjoint expression is indicated in red (negative COEX values). **C, D** and **E**. Different dimensionality reduction plots (Principal Component Analysis, t-distributed Stochastic Neighbour Embedding and Multidimensional Scaling, respectively) of 1234 genes (10% highest localized GDI). The primary markers used to set the variables are labeled. Dot colors denote the gene cluster corresponding to a layer or progenitor gene as specified in Section 2.5. Labels corresponding to other known layer identity markers (*Tbr1, Mef2c, Nes* and *Sox2*) that were not used as primary markers are also shown as additional landmarks. All plots refer to E17.5 mouse cortex cells^20^.

When GPA is performed genome-wide, it is possible to score genes according to how often their expression shows dependence on the expression of other genes, identifying cell-identity genes. To this purpose, we define the global differentiation index (GDI) of a gene *g* as a normalization of the average of the 5% lowest *p*-values obtained by GPA tests of *g* against all other genes.

We tested our method on neural development datasets. Indeed, brain embryonic structures display high cell diversity, with dividing multipotent progenitor cells, newborn neurons differentiating with many distinct identities and glial cells co-existing in a mixed cell population. This feature makes them particularly suited for scRNA-seq studies aiming to depict cell state identity and relationships between gene expression. We assayed COTAN on a scRNA-seq dataset of embryonic hippocampus^16^, focusing on a collection of selected constitutive genes (CGs^17^), neural progenitor genes (NPGs^18, 19^) and pan-neuronal genes (PNGs^18, 19^). COTAN’s GPA effectively discriminated between CGs, showing COEX near zero against all genes, and NPGs or PNGs, having positive or negative COEX when tested against one another (Figure 1A). Notably, each NPG positively correlated with other NPGs and negatively with PNGs, and vice-versa, indicating COTAN capability to correctly infer joint or disjoint expression of two genes at single cell level.

We compared COEX to Pearson correlation computing gene expression levels by Seurat^15^. COEX proved more accurate in discriminating between CGs, NPGs and PNGs, indicating COTAN as better suited in analyzing the co-expression of couples of genes at single cell level (compare Figure 1A with B). Notably, some CGs (e.g. *Calm1, Actb, Gapdh*) showed specific, high co-expression or disjoint expression with NPGs or PNGs when analyzed with Pearson correlation (Figure 1B), making them false positive markers of differentiated cell identity. We then compared GDI to the highly variable feature analysis of Seurat^15^. GDI efficiently discriminates between CGs, which lay below the median (with two exceptions, *Taf1* and *Zfr*), and NPGs and PNGs, located above the third quartile (Figure 1C). Instead, highly variable features analysis of Seurat (Figure 1D) was much less precise in discriminating between CGs and cell identity genes (compare Figure 1D to 1C).

Correlation analysis approaches are commonly used to identify cell clusters with consistent global gene expression. Conversely, we used COTAN to determine clusters of genes characterized by their co-expression patterns with selected genes in single cell. We chose to investigate mouse embryonic cortex^20^ because the molecular identity of a number of layer-specific precursor cells and neurons is known^18^. We firstly chose from literature^18^ robust primary markers for layer I (*Reln*), layers II/III (*Satb2*), layer IV (*Sox5*) and layer V/VI (*Bcl11b*), see Figure 2A. Then, for each marker we selected the most positively associated genes (COEX > 0 and GPA *p*-value < 0.0001 for *Satb2, Reln* and *Sox5* or < 0.001 for *Bcl11b*, for a total of 248 genes). For all these genes, we plotted an ordered symmetric heatmap of GPA COEX values with the genes grouped by the marker by which they were selected (Figure 2B). COTAN showed to be well suited to evaluate the co-expression of gene pairs genome-wide. The comparison between groups highlighted impressive consistency of co-expression inside each group and robust disjoint expression between different groups, with the only exception of *Sox5* and *Bcl11b* groups, which resulted as co-expressed. We believe that the *Reln, Satb2* and *Sox5* /*Bcl11b* groups represent genuine gene signatures of distinct cortical cell identity and that similar signatures can be found by unbiased approaches.

Eventually, we assayed the ability of our approach to cluster groups of co-expressed genes regarding cell layer identity (Figure 2C-E). In analogy to other methods^22^ the analysis was guided by few key genes. We chose ten primary layer identity markers (Greig at al 2013^18^): *Reln* and *Lhx5* for layer I (dark red), *Satb2* and *Cux1* for layer II/III (pink), *Rorb* and *Sox5* for layer IV (pale blue), *Fezf2* and *Bcl11b* for layer V/VI (dark blue), *Hes1* and *Vim* for neural progenitors (cyan). We then selected for each primary marker the 25 genes with the lowest GPA p-values. In this way we collected a total of 215 secondary markers (35 genes were shared by two primary markers). These were used as variables/features for the dimensionality reduction and cluster analysis. A normalization of COEX served as input data for the analysis (for details see Section 2.3 and 2.4). Figures 2C-E show PCA, t-SNE and MDS, respectively, of the 10% genes with highest localized GDI (*n* = 1234). The dots are colored according to a hierarchical clustering on the same input data (Section 2.5). Notably, each cluster shows univocal correspondence with all the primary markers of one of the major cortical cell identities at the developmental stage of analysis, proving COTAN ability to gather genes with similar nature regarding cell identity.

Eventually, this analysis highlighted a larger number of novel markers of cortical layer identity compared to the ones identified by conventional analysis.

### 2.2 Negative and synthetic dataset analysis

COTAN is a tool to study cell differentiation in scRNA-seq datasets. As such, it is important to assess its behaviour under the null hypothesis that there is no differentiation at all, i.e. on datasets that happen to be completely homogeneous. We refer to genes as constitutive if they are assumed to be transcribed in all cells of a population: their UMI counts may still be zero in some cells, but that would be because of random sampling. Therefore for homogeneous datasets all genes should be constitutive, while for normal multiple cell-type populations, only some of them are. For embryonic cortex datasets in this paper a number of constitutive genes selected from literature^17^ were included in our analysis.

We tested COTAN on three different types of homogeneous datasets: 1) a technical negative dataset formed by External RNA Controls Consortium mRNA (ERCC^7^), 2) a biological negative dataset formed by FACS sorted CD14+ cells^7^ and 3) two synthetic datasets with a single cell type (see Section 3.5).

We analyzed GPA and GDI on these datasets. Figure 3 shows that GPA’s *p*-value is a good approximation for the true *p*-value of the independence test. If the test is formally performed as usual by comparing the *p*-values with some given significance level, the false positives can be controlled efficiently.

**Figure 3.**
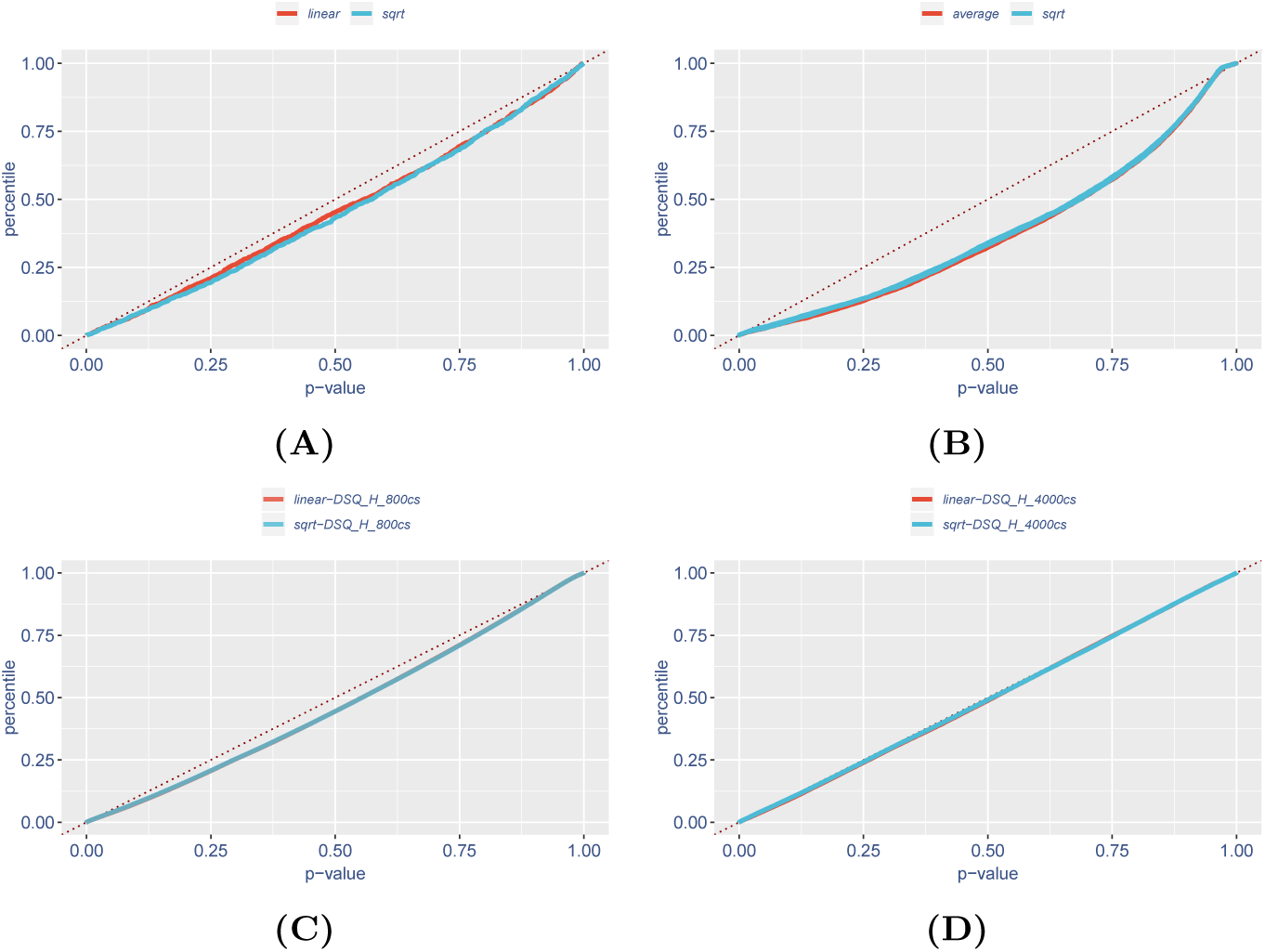
Empirical cumulative distribution functions of negative datasets GPA *p*-value. Points are a random sample from all 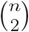 gene pairs. The plots are close to the identity function, so this statistics approximates a rigorous *p*-value. The fraction of expected false positives at significance level *α* is about *F* (*α*). Since deviations from identity are negative, false positive are *α* or less. **A**. ERCC mRNA dataset formed by 92 different mRNAs and 1015 droplets^7^; **B**. CD14+ cell dataset formed by 11405 genes (before cleaning) and 2370 cells^7^; **C, D**. Homogeneous synthetic datasets with 800 or 4000 cells.

COTAN’s GDI, applied on negative datasets (Figure 4), shows no unwanted dependence from total gene expression, and only few genes exceed the empirical threshold value 1.5 for differential expression, giving less than 1% of false positive.

**Figure 4.**
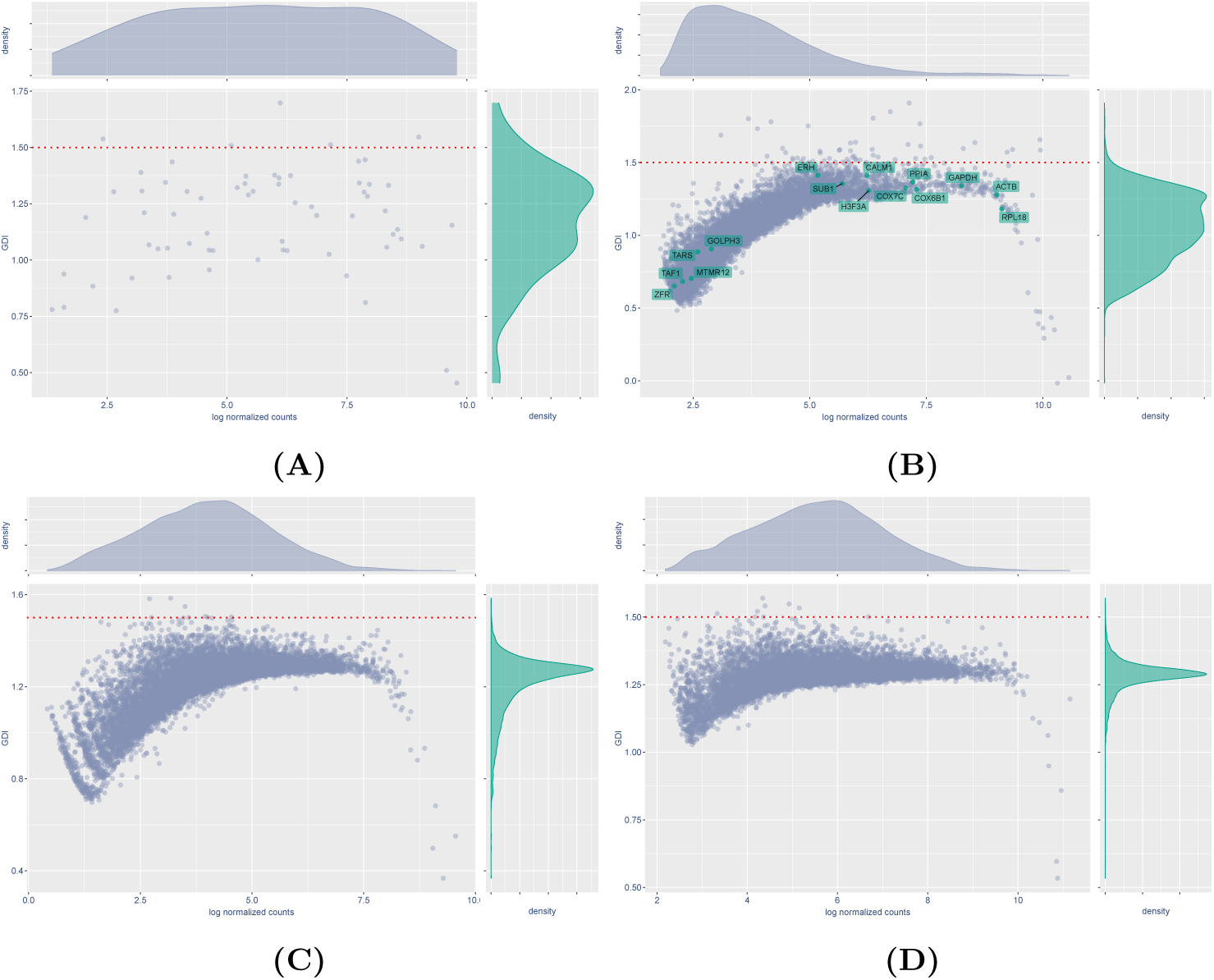
Negative dataset GDI. Plots show GDI vs log-normalized total UMI counts. High values of GDI are not more frequent for high or low UMI counts: GDI effectively keeps false positives controlled. The red dotted line is the empirical threshold 1.5 for differentially expressed genes. **A**. ERCC mRNAs; only 61 of the original 92 genes can be used in COTAN, as the other have non-zero UMI counts in all cells; there are only a few genes with GDI above 1.5. **B**. CD14+ cells; a total of 53 genes have GDI > 1.5, and they show enrichment for dendritic cell markers (Figure 5); the labelled sample of known constitutive genes shows low GDI values both with strong and weak expression. **C, D**. Homogeneous synthetic datasets with 800 or 4000 cells; there are only a few false positive genes with GDI above 1.5.

We noticed that CD14+ cell dataset exhibits some high GDI genes (Figure 4B) and decided to investigate their nature. Their COEX heatmap (Figure 5) displays both positive and negative values, hint of the presence of at least two cell subtypes inside this nearly uniform dataset. In fact we found that the set of these high GDI genes is enriched in dendritic cell markers, as indicated by gene enriched analysis performed by Enrichr^23, 24^ web site. The analysis found a subset of 38 enriched genes, with adjusted *p*-value 2.32 · 10^−22^ regarding ARCHS4 Tissues database. As reported also in the original paper^7^, a small sub-population of this cell type was in fact detected in the CD14+ cell dataset, in agreement with our findings.

**Figure 5.**
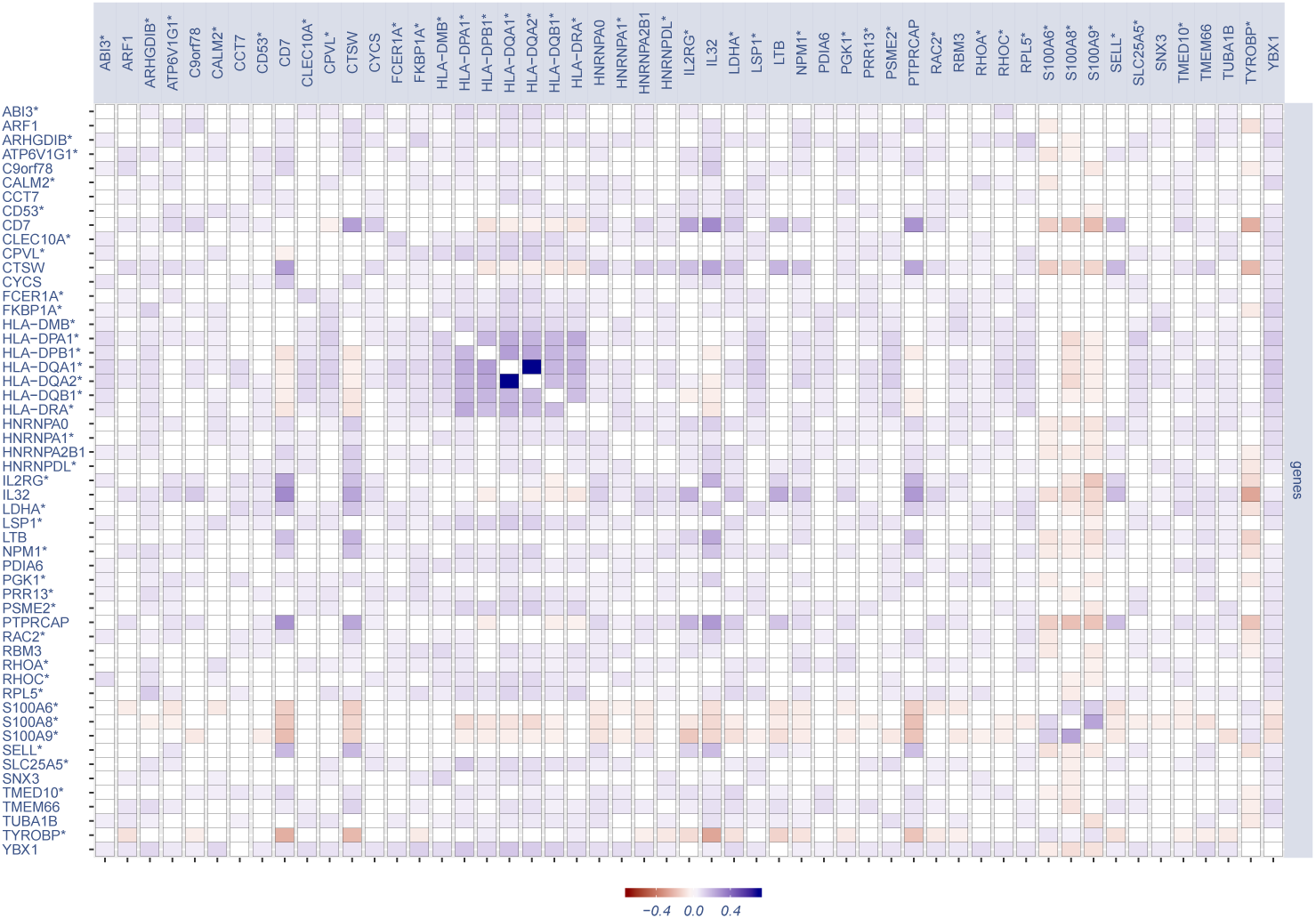
GPA COEX heatmap for the 53 genes with GDI greater than 1.5 in the CD14+ cell negative biological dataset. Most genes associated by positive co-expression are coherent with the existence of a dendritic cell subpopulation, and therefore their high GDI does not represent a false positive. Genes marked with * are those found by Enrichr (see text).

### 2.3 Gene co-expression space plots

Because COTAN was specifically developed to analyze joint or disjoint expression of two genes, it is possible to analyze which genes present similar behaviour in the dataset. The basic idea is to select a set *V* of informative genes (as variables/features) and then use COTAN to represent all genes, genome wide, according to their pattern of co-expression with respect to those in *V*. This multivariate representation allows then to cluster genes by their co-expression pattern and, after a dimensionality reduction, to plot them on the plane.

We chose a comprehensive set *V* of layer-associated genes to be used as variables (Figure 6). To build *V* we selected a set of ten known primary markers of cortical layer identity (*Reln* and*Lhx5* for layer 1, *Satb2* and *Cux1* for layer 2/3, *Rorb* and *Sox5* for layer 4, *Bcl11b* and *Fexf2* for layer 5/6 and *Vim* and *Hes1* for progenitor cells), together with the top 25 genes most correlated with each of them, for a total of 215 secondary markers (this number is lower than 250 because some of these genes were shared).

**Figure 6.**
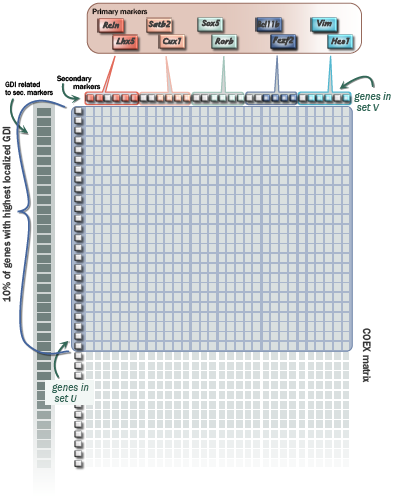
Schematic of the procedure to build the sets of genes *V* and *U* used for dimensionality reduction analyzes (see text).

For all genes in the dataset, we computed the *localized GDI* relative to the genes in *V* (see Section 2.4 below) and used it to filter only the 10% mostly differentially expressed genes, with the highest values, in order to get a meaningful graphical representation and better input data for the subsequent cluster analysis (Figure 6). These selected genes formed the set *U* of units/points to represent.

Then, as described in Section 3.6, we filled the matrix *U* × *V*, applied the normalization tanh 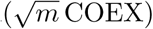, yielding the gene co-expression space, and finally performed three different dimensionality reduction analyzes on the resulting index: PCA (Figure 2C), t-SNE (Figure 2D) and MDS (Figure 2E). t-SNE plot was performed using 20 initial dimensions, 3000 maximum iterations, perplexity 30, eta = 200 and theta = 0.4. Genes were colored according to the cluster analysis results (see Section 2.5).

### 2.4 Localized GDI

Global differentiation index (GDI) as defined in Section 3.4 is a method to score genes according to their propensity to show statistically significant joint or disjoint expression against *all* other genes. Although this is not a rigorous test of differential expression, we remark that values around 1.3 are typical for average-expression, constitutive genes, while values above 1.5 are evidence of differential expression. To make this tool more specific, it is possible to restrict the set of genes against which GDI is computed to a selected subset *V*, with the recommendation to include a consistent fraction of cell-identity genes (for instance markers of layer identity). In this case the index is referred to as GDI localized to *V*.

Localized GDI may uncover information that is hidden or confounded in the genome-wide GDI. We stress that the differentiation threshold 1.5 is referred to the latter, and that the former should be used only to score genes and not to infer whether they are in fact constitutive for the cell population or not (Figure 7).

**Figure 7.**
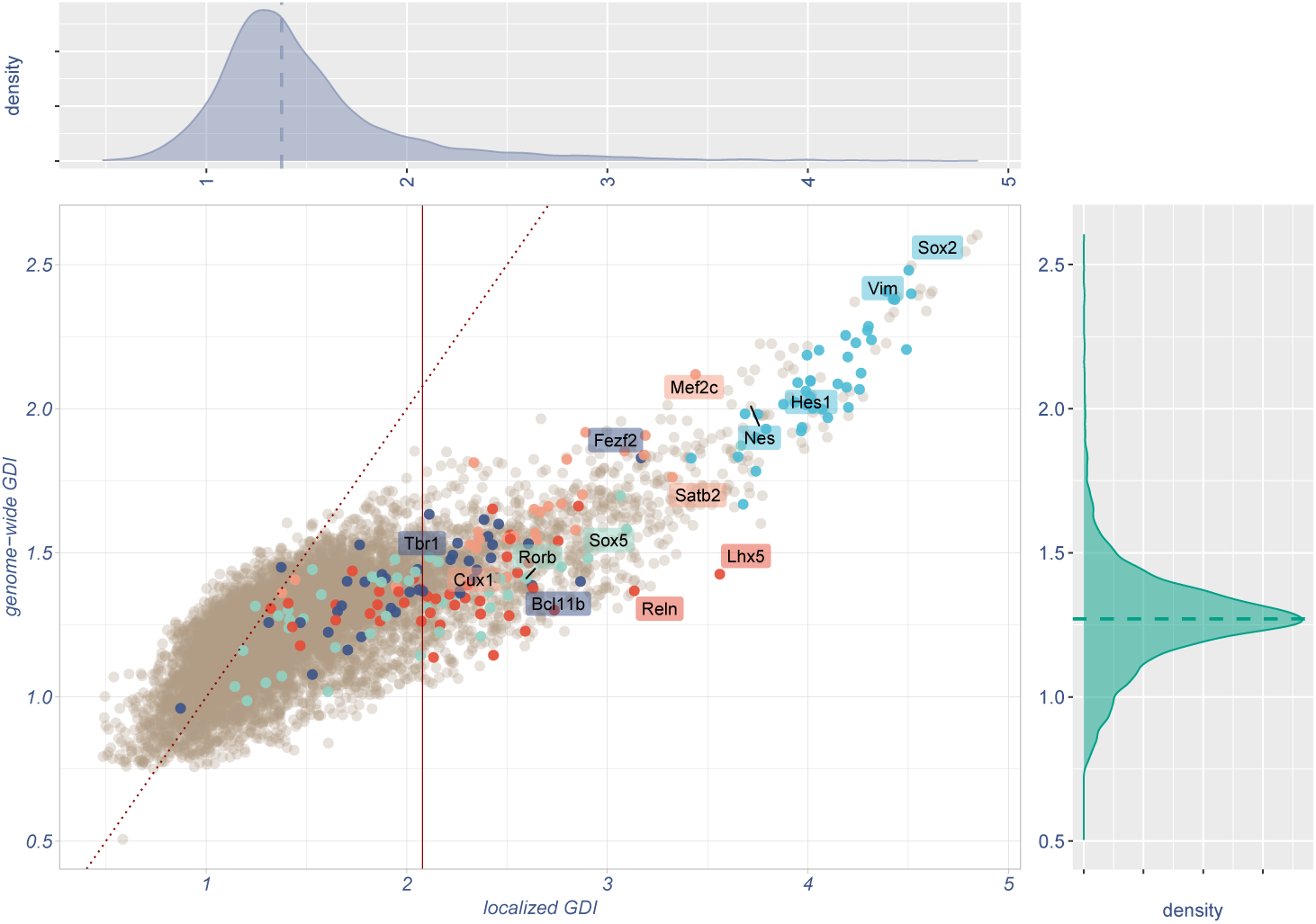
Comparison between genome-wide and localized GDI for all genes of E17.5 mouse cortex^20^. Labels and respective colors as in Figure 2, in particular only primary markers selected for layer identity together with landmark layer identity genes are labelled. Colored dots are those secondary markers (i.e. the set *V* of Section 2.3) that have positive COEX against a primary marker, with which it shares the color. Dotted red line is the identity function. Localized GDI has typically higher values than global GDI, and its right tail is more spread. Notably, many informative genes of layer identity have intermediate global GDI, below the differentiation threshold 1.5. It would be difficult to detect them without localized GDI. The vertical line corresponds to the 90-percentile of the localized GDI and its right side identifies the 10% genes used for cluster analysis.

### 2.5 Gene clustering

We performed a cluster analysis on the 10% genes selected in Section 2.3, using their co-expression space representation (see Section 3.6) as an input for a hierarchical clustering with the Ward’s method. The resulting tree presents a natural cutting distance at seven clusters (possible alternatives being at 2, 3 or 5 clusters – see Figure 8).

**Figure 8.**
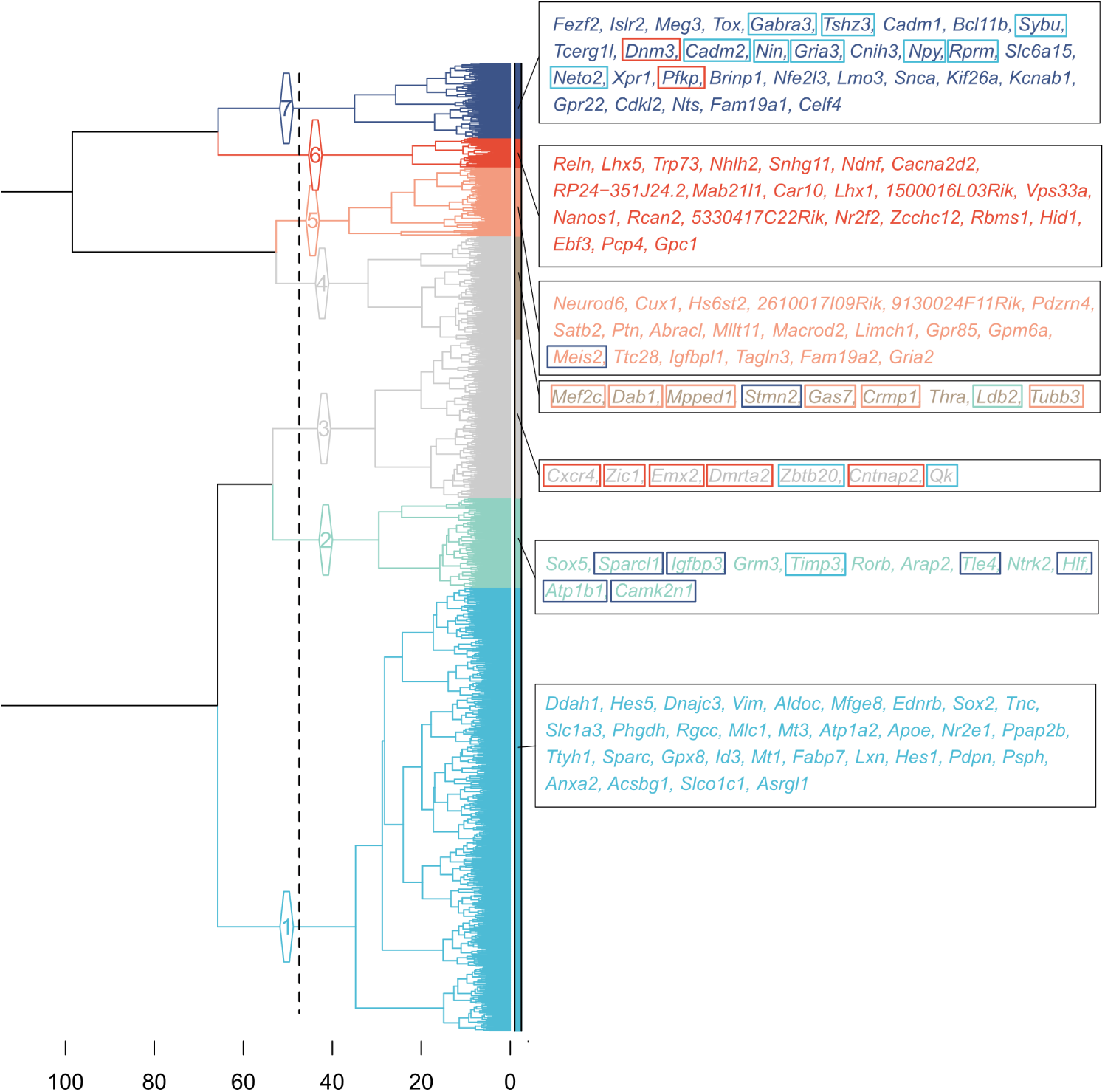
Hierarchical clustering of genes. The dotted line denotes the height of the tree cut forming seven clusters. Branches and leaves colors indicate cluster identity: cluster 1, in cyan, indicates progenitor identity, cluster 2, in light blue, indicates layer IV identity, cluster 5, in pink, denotes layer II/III identity, cluster 6, in dark red, indicates layer I identity and cluster 7, in dark blue, indicates layer V/VI identity. The two clusters in gray (3 and 4) do not contain primary markers and are likely inconsistent with projection neuron identity (see text). The gene names reported are the ones identified as secondary markers (see Section 2.3); among these, the ones initially linked to a primary marker not coherent with the final cluster are boxed with the color related to their initial marker.

In five of the seven clusters we found the five pairs of primary markers, each one in its own. From them we assigned the identity of the five clusters and in particular: cluster 1, containing *Vim* and *Hes1*, was identified as genes related to progenitors identity, cluster 2, containing *Sox5* and *Rorb*, was identified as genes related to layer IV identity, cluster 5, containing *Cux1* and *Satb2*, was identified as genes related to layer II/III identity, cluster 6, containing *Reln* and *Lhx5*, was identified as genes related to layer I identity and, finally, cluster 7, containing *Bcl11b* and *Fezf2*, was identified as genes related to layer V/VI identity.

To asses the ability of COTAN gene clustering to detect new markers we compared our clusters with the literature^25^. Results are summarized in Tables 1 and 2.

**Table 1.** Number of layer markers found by Loo et al.^25^ with their respective layer according to the original paper (columns) and according to COTAN (rows). Bold text denotes consistent identification by the two methods. *Plxna4* in purple, see text.

|  |  | Markers from Loo et al.<sup>25</sup> |  |  | Prog |
| --- | --- | --- | --- | --- | --- |
|  |  | Layer I | Layer II/III | Layer V/VI |  |
| Markers<br>detected by<br>COTAN | Layer I | **4** |  |  |  |
|  | Layer II/III |  | **2** | **1** |  |
|  | Layer V/VI |  |  | **9** |  |
|  | Prog |  |  |  | **7** |
|  | Other cluster |  |  | 2 | 1 |

**Table 2.** Same as Table 1, but excluding all genes belonging to the set *V* of secondary markers, as these were selected by co-expression with the primary markers and hence their assignment to the correct clusters might be favored by the method.

|  |  | Markers from Loo et al.<sup>25</sup> |  |  | Prog |
| --- | --- | --- | --- | --- | --- |
|  |  | Layer I | Layer II/III | Layer V/VI |  |
| Markers<br>detected by<br>COTAN | Layer I |  |  |  |  |
|  | Layer II/III |  | **1** | **1** |  |
|  | Layer V/VI |  |  | **5** |  |
|  | Prog |  |  |  | **4** |
|  | Other cluster |  |  | 2 | 1 |

We can observe from the tables that all markers used or identified by Loo et al.^25^ are also detected by COTAN clustering and the agreement in the layer assignment is quite good. There are only 4 genes out of 26 that have been assigned to different layers. *Plxna4*, which is the only one with known cortical expression pattern in Allen Brain Atlas database, is recognized by Loo et al. as marker for layer V/VI at stage E14.5, and by our analysis as marker for layer II/III, at stage E17.5. Accordingly, ISH Allen Brain Atlas in Figure 9 shows that this protein is actually localized principally in layer V/VI at early stages, but it co-localizes with layer II/III at later stages.

**Figure 9.**
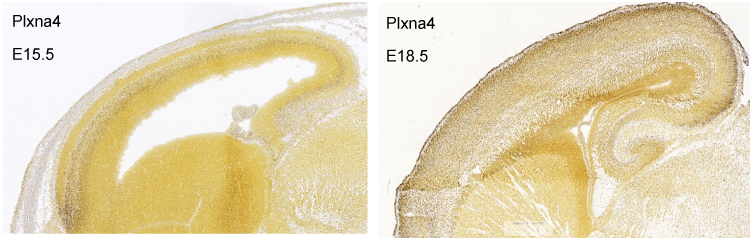
In Situ Hybridizations data from Allen Brain Atlas website^26^ for mouse cortex at E15.5 and E18.5. It is evident that *Plxna4* protein at early stage is principally localized in layers V/VI, while at later stages it is more present in superficial layers, in accordance with both our finding and the work from Loo et al.^25^

Notably, COTAN identified a much higher number of layer markers compared to the conventional methods applied by Loo et al. (see supplementary file `STable1.csv`). Among all possible new layer markers detected by COTAN, we analysed the ones presenting nucleic acid binding gene ontology (GO:0003676). Complete tables are attached as supplementary files (`STable1.csv and STable2.csv`). Figure 10 shows the E18.5 ISH collection of the genes available from Allen Brain Atlas website. Most of the genes show ISH pattern consistent with layer identity as identified by COTAN, with few exceptions.

**Figure 10.**
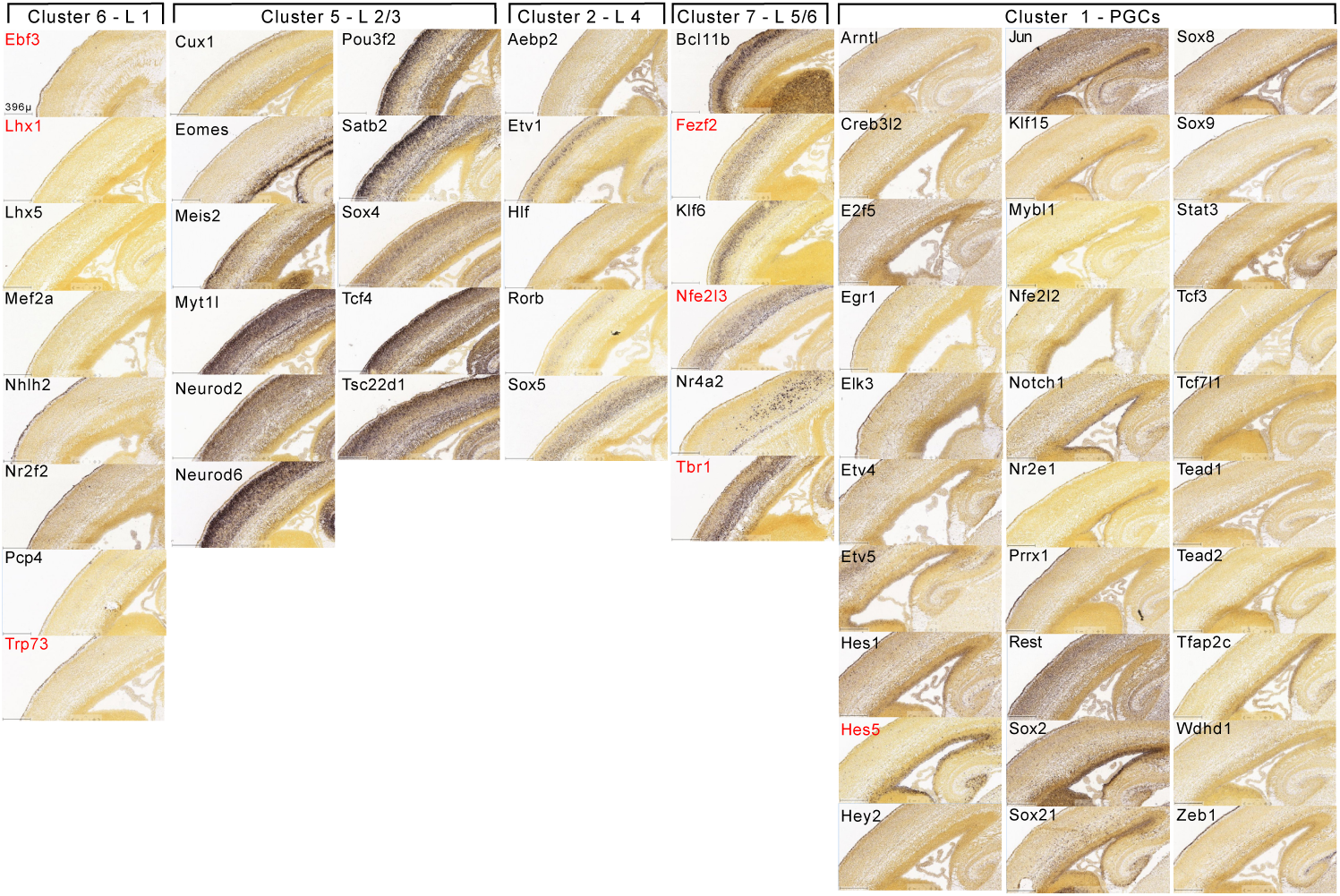
E18.5 ISH from Allen Brain Atlas website^26^ of markers with nucleic acid binding gene ontology identified by COTAN. In red, markers detected also by conventional analysis are indicated.

Among the seven natural clusters detected by hierarchical clustering, two clusters do not contains any primary markers. To assess their identity we performed a gene enrichment analysis on the Enrichr web site^23, 24^. We found that cluster 3 is enriched in septofimbrial nucleus genes, and cluster 4 is enriched in accumbens nucleus genes (in the Allen Brain Atlas upregulated genes dataset – data tables attached as supplementary files).

## 3 Methods

### 3.1 Mathematical framework

We refer for all the details to the dedicated paper^14^. We resume here the main aspects of the mathematical framework.

#### 3.1.1 Model and parameters estimation

For each gene *g* and cell *c*, let *R* denote the UMI count and 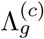 the corresponding biological expression^1^. Conditional on the value of the latter, the read count has 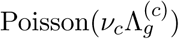 distribution, where *ν*_*c*_ denotes the UMI detection efficiency (UDE) of cell *c*, which is not supposed to depend on the gene (the pipeline in Section 3.7 includes a step to check this assumption). Then the expected UMI count is given by 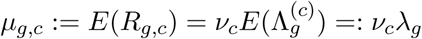. We introduce two different frameworks for estimating these parameters: *linear* and *square-root*.

With the symbol ∗ used to denote averaging along all the values of the corresponding index, and under the convenient assumption that *ν*_∗_ = 1, then *R*_*g*,∗_, *R*_∗,∗_ and *R*_∗,*c*_ are unbiased estimators of *λ*_*g*_, *λ*_∗_ and *ν*_*c*_*λ*_∗_, and 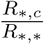, is an estimator of *ν*_*c*_. These are called the *linear estimators* and may often be used with good results, though sometimes they may not be very robust, when there is a large biological variability of the highly expressed genes. (In particular, the estimator of *ν*_*c*_ may end up being correlated with the expression of these genes.)

A possible alternative is using the variance-stabilizing transformation 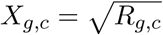 which yields the *square-root estimators*, defined as follows:

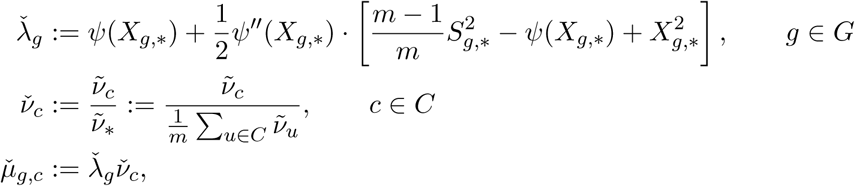

where *ψ* is the inverse of the function 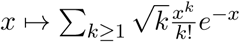 on [0, ∞), where 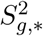 denotes the sample variance of *X*_*g,c*_ for fixed *g*, and where the “tilde” estimator of *ν* is the analogous for cells of the “check” estimator for *λ*, so:

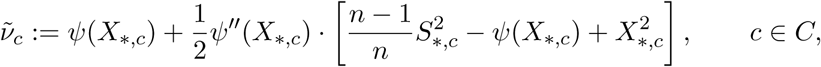

We remark that *ψ*(*x*) ≈ *x*^2^ and in particular the above formulas are just a second order correction to the naive attempt at commuting of the square root and the average.

Accuracy and precision of these estimators were tested on synthetic datasets with heterogeneous cell types, for which the true values of *ν* and *λ* were known (Figure 11 and Section 3.5).

**Figure 11.**
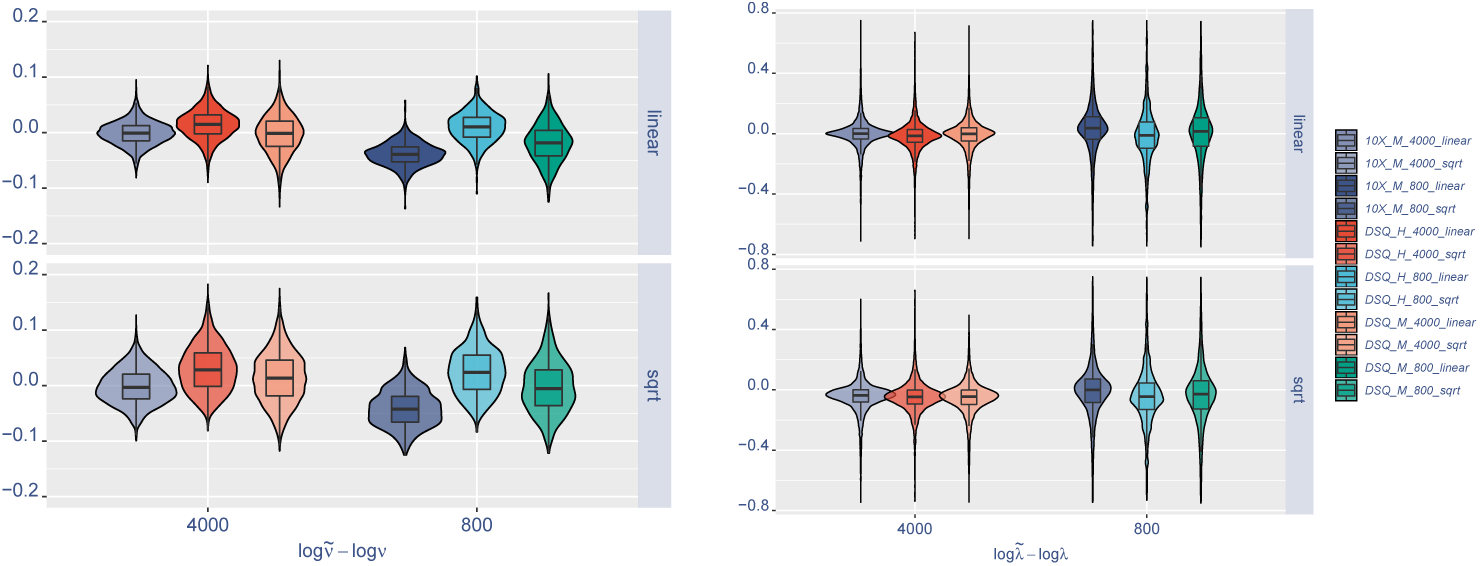
Violin plots of linear and square-root estimator error. The six synthetic datasets differ for the number of cell types and the total number of cells. Homogeneous datasets are for reference. In the legend `10X` and `DSQ` (Drop-seq) refer to the method of the real dataset used as a source. `H` and `M` indicate if the dataset is homogeneous or with multiple cell types; `800` and `4000` refer to the number of cells simulated, and `sqrt` and `linear` indicate the type of estimation used for *ν* and *λ*.

#### 3.1.2 Probability of zero UMI count

Whichever the estimator of *µ*_*g,c*_ = *λ*_*g*_*ν*_*c*_, its value is the starting point to approximate the probability that *R*_*g,c*_ = 0. In fact by the Poisson conditional distribution of the UMI counts, we get

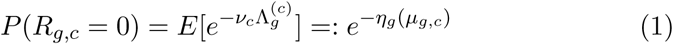

where *η*_*g*_ denotes the log-mgf function of 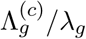. We don’t fix a partic-ular model for the distribution of either 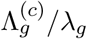 or *R*_*g,c*_, but we make the assumption that at least the probability mass in zero for *R*_*g,c*_ can be approximated by a negative binomial model with mean *µ*_*g,c*_ and some dispersion *a*_*g*_ depending only on the gene. Then equation (1) becomes

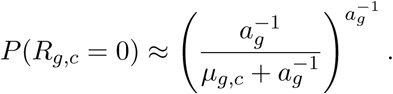

The value of *a*_*g*_ is estimated by imposing the marginal condition that the expected number of cells with zero UMI count for the gene, 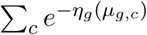, equals their total number: with # denoting cardinality,

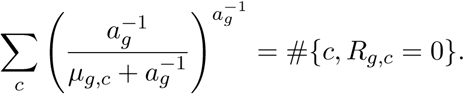

In the case of the genes that would require a negative dispersion, because 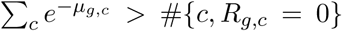, we choose instead to impose a zero dispersion model (Poisson distribution) with *increased mean* (1 + *b*_*g*_)λ_*g*_, yielding 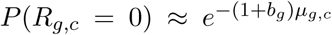. This choice is consistent with the intended universality of the approach, because no distribution of 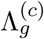 would give a negative dispersion and because *λ*_*g*_ is anyway an estimated quantity and therefore noisy. In the typical dataset about 30% of all genes fall in this case, with values of *b*_*g*_ no larger than 0.02 in average and with no single value higher than 0.15. In the implementation, *a*_*g*_ and *b*_*g*_ are encoded in a single parameter (Section 3.7.3).

### 3.2 Co-expression tables

COTAN at its core is a tool for measuring co-expression of two genes. In this section we take advantage of the estimators introduced above, and show how to create and test the co-expression tables on which COTAN’s *gene pair analysis* (GPA) is based.

Gene co-expression is a powerful tool to study cell differentiation^27^. When the population of cells is completely homogeneous, each gene is assumed to be either expressed in all cells or not expressed in any. In this situation the UMI counts of the expressed genes should be independent random variables across the cells. On the other hand, if the population is non-homogeneous and there are different cell types, each one expressing its own genes, then two genes could have non-zero UMI counts in the same cells, more (or less) often that should be expected if the population was homogeneous.

COTAN is built on the biological assumption that cell differentiation will typically *shun to zero* the expression of several genes, and that UMI count fold change is not as much informative as the switch of a gene from present to absent (or vice versa). In fact fold change is also difficult to assess for genes with very low expression, which are the most common in scRNA-seq. Significantly, in these datasets several constitutive genes have very few UMI counts (Figure 12). However, being constitutive, their low counts cannot be considered indication for absent or negligible gene expression. Consequently, for low-expression genes the main information is whether their UMI count is positive or zero.

**Figure 12.**
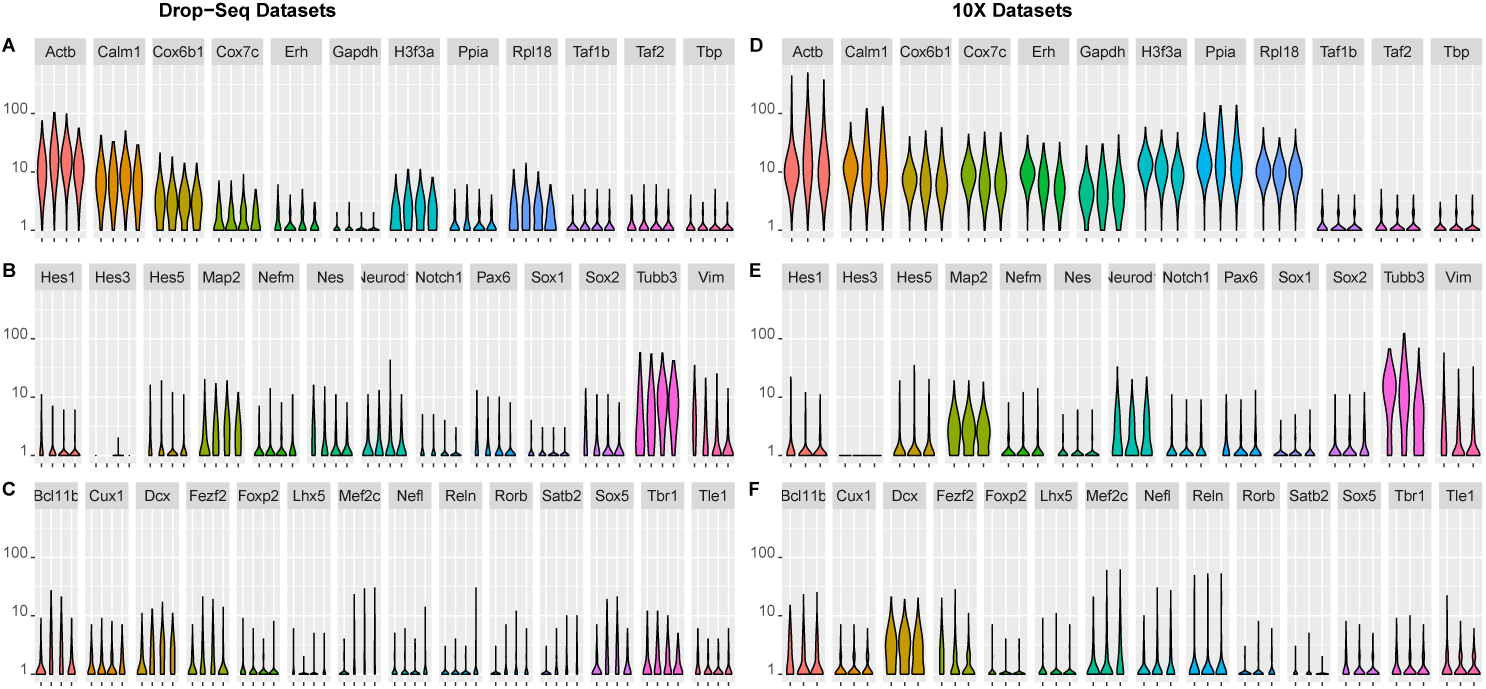
Violin plots of UMI counts + 1, with *y*-axis in logarithmic scale. On the left column (**A, B, C**) plots of a Drop-seq mouse cortex dataset^20^ at four development times (E11.5, E13.5, E15.5 and E17.5). On the right column (**D, E, F**) plots of a 10X dataset^16^ showing the development of mouse dentate gyrus (E16.5, P0 and P5). **A, D**. Constitutive genes. **B, E**. Selected neural markers. **C, F**. Cell identity transcription factors. Notably, the UMI counts of some constitutive genes, such as *Taf1b, Taf2, Tbp* and *Gapdh*, are quite low. Neural markers and identity genes with similar UMI count levels, such as *Sox1, Notch1, Tle1* or *Satb2*, are expected to be expressed with biological relevance.

From a mathematical point of view, moreover, the very low typical expression of most genes is also the main cause of dropout artefacts and more in general of the difficulties in modelling gene expression levels^8, 9^. In fact log-normalization, even with pseudocounts, essentially assumes to act on larger numbers, so that they may be thought of as continuous variables, but scRNA-seq datasets exhibit instead discrete counts, typically with 95% counts of 2 or less and 98% of 4 or less.

Based on the previous biological assumption, and on the mathematical need to deal with the discrete UMI count variable, we choose to collapse the values *R*_*g,c*_ in the two categories {*R*_*g,c*_ = 0} and {*R*_*g,c*_ ≥ 1}, neglecting to consider values larger than 1 as different. COTAN GPA main test compares the number of cells with zero read count in couples of genes (jointly versus marginally), in a way similar to 2 × 2 contingency tables, but generalized to experimental units with different UDE.

For example consider the constitutive genes *Cox7c* and *Tars* in the following case:

| *Cox7c* vs *Tars* | $Tars \geq 1$ | $Tars = 0$ | tot |
| --- | --- | --- | --- |
| $Cox7c \geq 1$ | 262 | 1006 | 1268 |
| $Cox7c = 0$ | 249 | 1376 | 1625 |
| tot | 511 | 2382 | 2893 |

This contingency table *cannot* be tested with the standard statistics, which are built either on the hypergeometric distribution (Fisher’s test) or on deviation from expected values given by the proportion of the marginals (chi-squared test). In fact, these methods are based on the assumption that all experimental units have the same probabilities of ending in the four counts. In scRNA-seq data, the different UDE *ν*_*c*_ of different cells makes this hypothesis false, and the standard test would give *spurious correlations*, as can be seen by computing the expected values in the standard way:

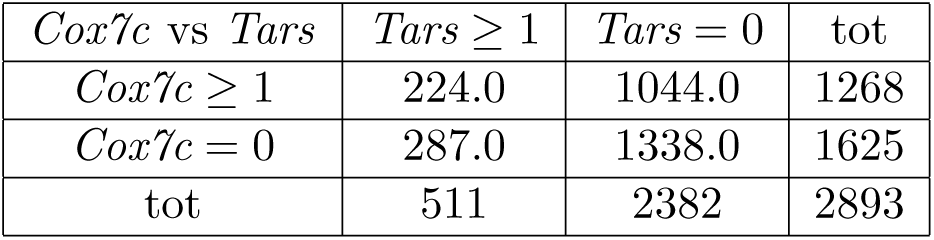

*Cox7c* and *Tars* are constitutive genes, and hence they should not show a significant co-expression, but with classical statistics the resulting *p*-value is very low (0.019% with chi-squared test and 0.022% with Fisher-Irwin test). In fact the cells with lower UDE have higher probability to end in the 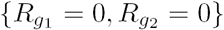 box and so they raise the frequency of that outcome above the expected 1338, yielding in this case a total of 1376 cells. With the classical contingency table interpretation, these would be explained by a positive co-expression of the two genes, but this is incorrect: the two genes are constitutive, and should be really expressed in all cells in the sample, it is random sampling that causes some cells to have zero UMI count. Random sampling of the two genes should also be independent, *conditionally* on the cell UDE, but if UDE is not taken into account, then there is correlation of the two genes *through UDE variability*.

Fortunately, with the estimates of *P* (*R*_*g,c*_ = 0) available, we can approximate the expected count of the 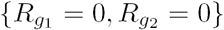 box with 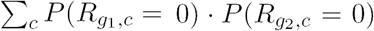. In the example above, this yields expected values which are much closer to the observed values:

| *Cox7c* vs *Tars* | *Tars* $\geq 1$ | *Tars* = 0 | tot |
| --- | --- | --- | --- |
| *Cox7c* $\geq 1$ | 256.1 | 1011.9 | 1268 |
| *Cox7c* = 0 | 254.9 | 1370.1 | 1625 |
| tot | 511 | 2382 | 2893 |

### 3.3 Test for GPA and co-expression index

The proposed GPA model yields an approximate table of expected cell counts under the hypothesis of independence of two genes *g*_1_ and *g*_2_, therefore it is possible to build the usual statistics 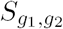 for contingency tables, by comparing observed counts *O*_*i,j*_ to expected counts *ϵ*_*i,j*_ (dependence on *g*_1_ and *g*_2_ implicit):

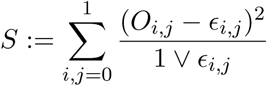

The small correction on the denominators is needed because we are forced to use this statistics also when some of the expected values are small, since Fisher’s test cannot deal with variable UDE’s. The correction is conservative, in the sense that it will not increase false positives.

One can then formally compute a *χ*^2^(1) *p*-value as usual as 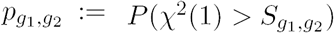. For example, in the case of the two constitutive genes of the previous section, one gets *p*_*Cox7c,Tars*_ = 56.4%, in accordance with the null hypothesis that the two genes are independently found in the cells.

Rigorously, the computed probability is not a true *p*-value for this test, since one cannot easily deduce its distribution under the null hypothesis, but it has the usual qualitative meaning, and tests on negative synthetic datasets (see Section 3.5) show it to be near to the uniform distribution on the unit interval (Figure 3C-D). Figure 13 shows the distribution of GPA’s *p*-values for several genes in a normal biological sample with different cell types^20^.

**Figure 13.**
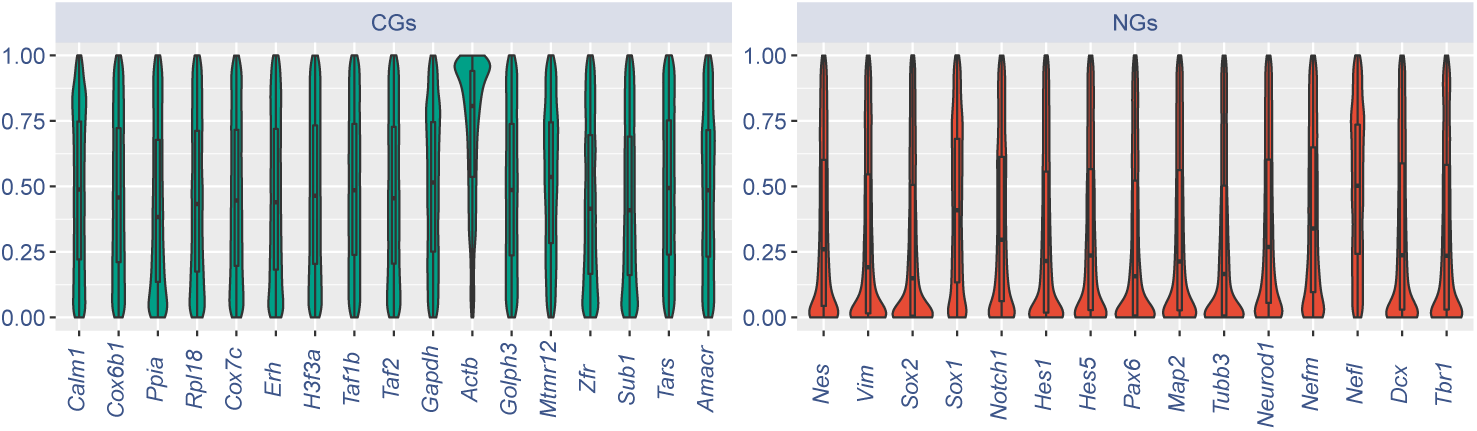
Violin plots of GPA’s *p*-values of CGs (left) and NGs (right), from^20^ E15.5. Each gene was tested for independence against all other genes and the resulting approximate *p*-values are represented. CG values are typically uniformly distributed, as expected (see Section 3.4). Small values are instead more frequent for NGs, as these are either co-expressed or disjointly expressed with most other non-constitutive genes.

We introduce also the GPA *co-expression index* (COEX) of the two genes, defined as 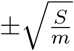, where *m* denotes the number of cells in the sample, and with the sign chosen according to whether the deviation of *O*_*i,j*_ from *ϵ*_*i,j*_ is in the direction of joint expression (positive) or disjoint expression (negative). Hence we could write 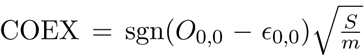 or other equivalent formula.

When the null hypothesis of the test is true, COEX distribution will be approximately Gaussian with mean zero and small variance 1*/m*, so values near zero are in accordance with the independence hypothesis. On the other hand, when the null hypothesis is false, COEX will converge, for *m* → ∞, to some deterministic coefficient in [−1, 1] depending on the joint expression of the two genes in the population, hence values away from 0 may indicate bonds between the genes^2^. Figures 1A and 2B show typical examples of COEX values for constitutive and cell-identity genes: it is apparent that COEX is a powerful tool for studying genes relations and cell differentiation.

### 3.4 Global differentiation index

When GPA is performed genome-wide, between all pairs of genes of the sample, it becomes possible to score genes based on their propensity to show either joint or disjoint expression against any other gene. Consider in particular a constitutive gene. It should be (biologically) present in all cells and its possible disappearance from the UMI counts of some cells should be only due to random sampling, independently from any other gene. Therefore, for any constitutive gene, the distribution of the GPA *p*-values against other genes should be approximately uniform on the unit interval (Figure 13, left). On the other hand, for a cell-identity gene, we expect a considerable fraction of *p*-values to be very small, as the independence hypothesis will be false for all the related cell-identity genes (Figure 13, right).

The global differentiation index (GDI) of a gene is defined as a normalization of the average of the smallest 5% of all the GPA *p*-values against other genes. The normalization chosen, *f* (*x*) = ln(− ln(*x*)), helps to make the index distribution more evenly spread and easy to grasp. Analysis of synthetic and negative datasets show that a value above 1.5 can be usually associated with non-constitutive genes, with < 1% false positives (Figure 4). For a constitutive gene with average expression level, a typical value for the GDI should be around ln(− ln(0.05*/*2)) ≈ 1.31. Observe that because the GDI comes from *p*-values, it is expected to depend on the sample size for non-constitutive genes: a larger cell number makes the statistical test more powerful producing GDIs generally higher, for differentially expressed genes.

### 3.5 Synthetic datasets creation

We generated six synthetic datasets to assess the performance of COTAN and the quality of the estimators.

To make them more realistic, the parameters were obtained from two biological datasets: an E17.5 mouse cortex Drop-seq (DSQ) dataset^20^ and a P0 hippocampal mouse 10X dataset^16^. For both of these, cells were clustered adapting the Louvain-Jaccard Clustering algorithm used in Loo et al.^25^, finding 6 and 15 cell types respectively. We assumed conditional 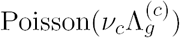 distributions for UMI counts, and for 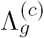 we assumed gamma distributions with parameters depending on cell type and on the gene. Then we performed maximum likelihood estimation on each cluster separately (with an iterative optimization procedure using Tensorflow^28^), to get the UDE *ν*_*c*_ for each cell and the gamma parameters for each cell type and for each gene.

After parameters estimation, the synthetic datasets were generated all in the same way (described below) but with different sources (DSQ or 10X), number of cells (800 or 4000) and cell types (1 or 6 for DSQ, or 15 for 10X). The two synthetic datasets with homogeneous (H) cell type (`DSQ_H_800` and `DSQ_H_4000`) are considered negative datasets and are used to assess the false positive rate of GPA and the distribution of approximated *p*-values and GDI under the hypothesis of independence (see Section 2.2). The four synthetic datasets with multiple (M) cell types (`DSQ_M_800, DSQ_M_4000, 10X_M_800` and `10X_M_4000`) are considered heterogeneous samples and are used to assess the performances of the parameters estimators (see Section 3.1.1).

For each dataset we generated the UDE of each cell *c* by sampling from the empirical distribution obtained with COTAN estimators on the biological samples (see Figure 16B-C for typical distributions).

**Figure 14.**
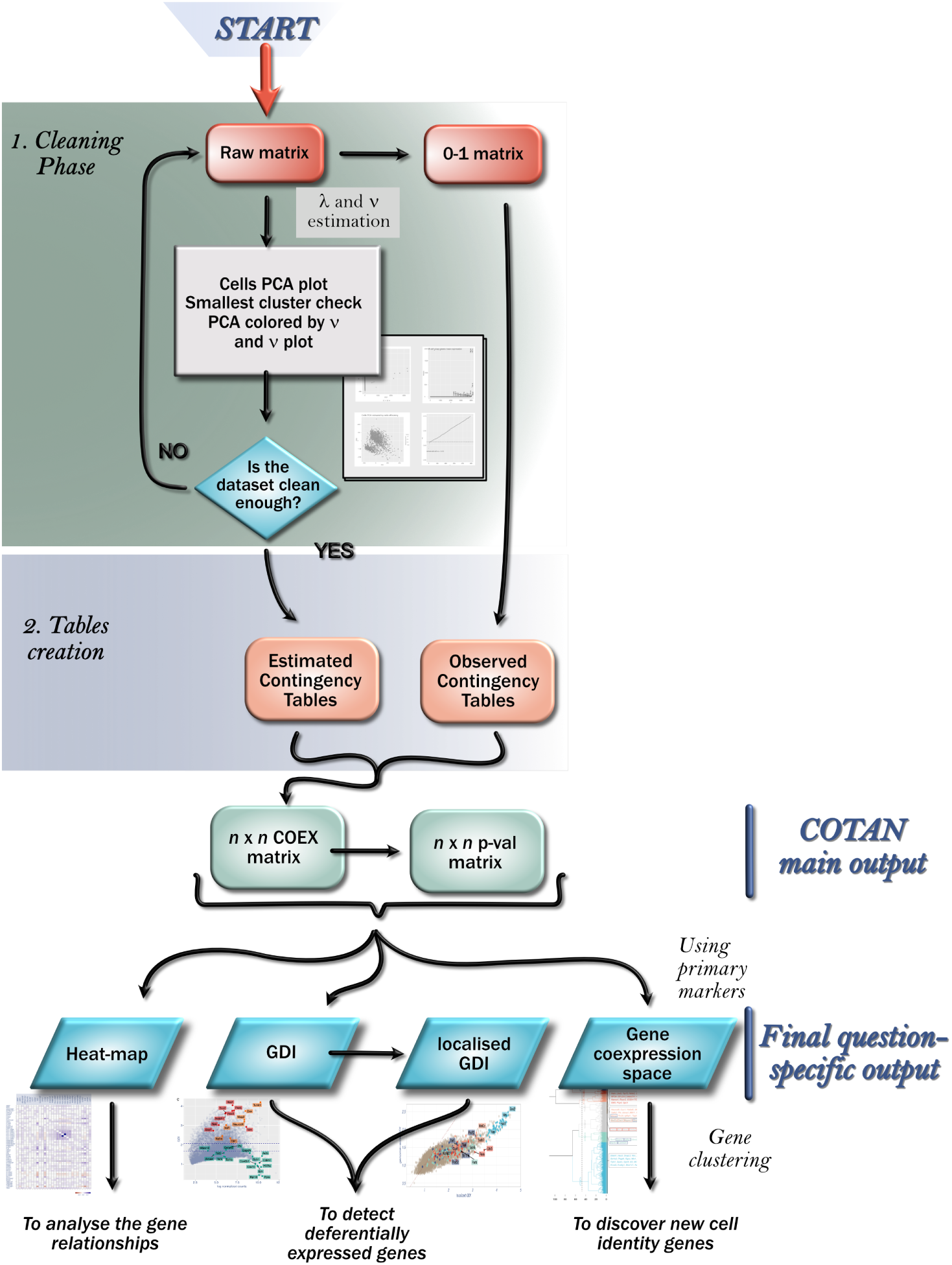
Pipeline diagram.

**Figure 15.**
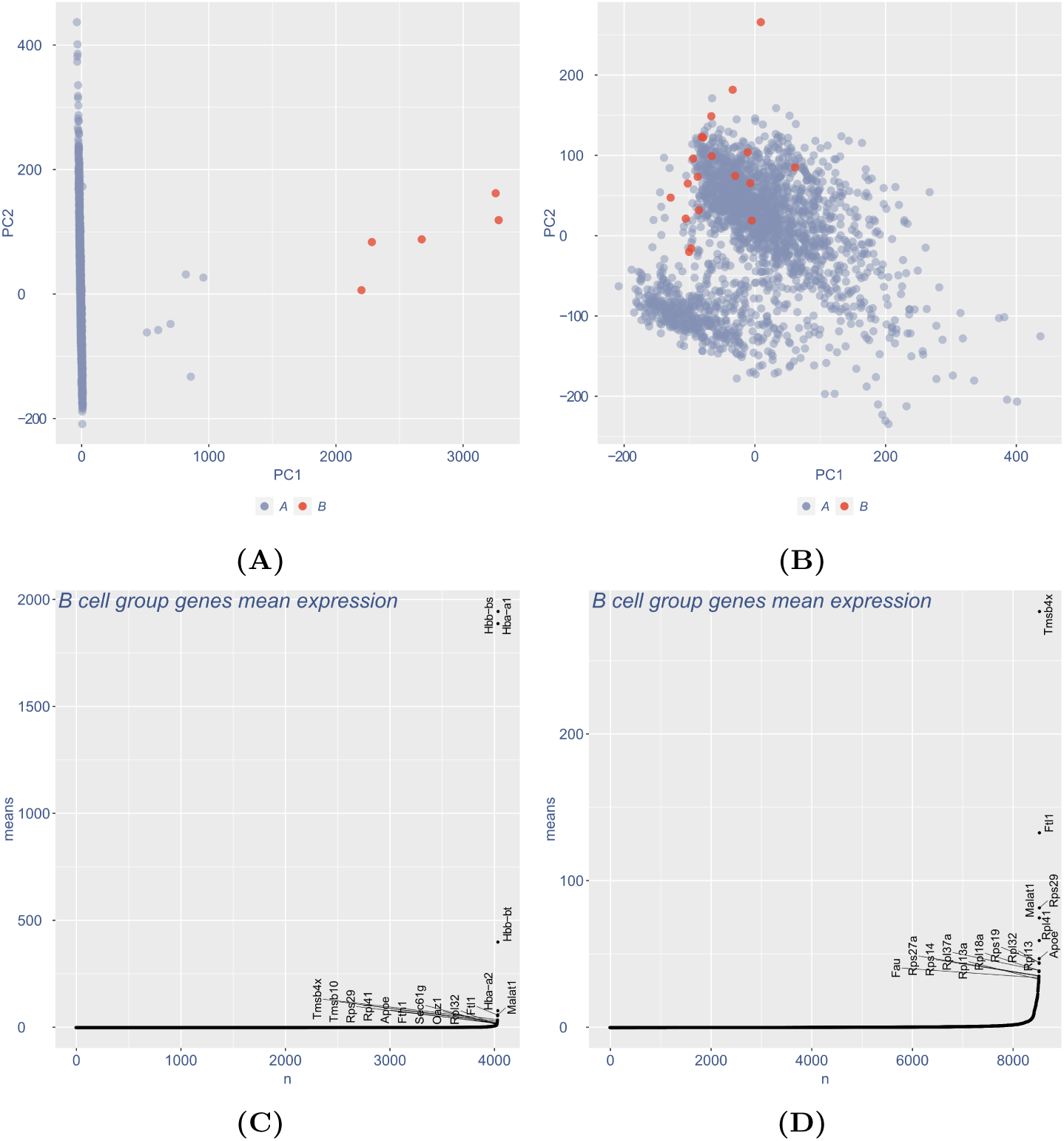
Cleaning plots. **A, B**. PCA plots of the cells clustered by Mahalanobis distance in a larger *A* (blue) and a smaller *B* (red – see text) set. **C, D**. Plots for the most expressed genes in the *B* cell group. **A, C** show a dataset that must be cleaned, with a small well separated cluster *B* and high expression on unusual genes. **B, D** show an example of the final stage, when no more cleaning iterations are required.

**Figure 16.**
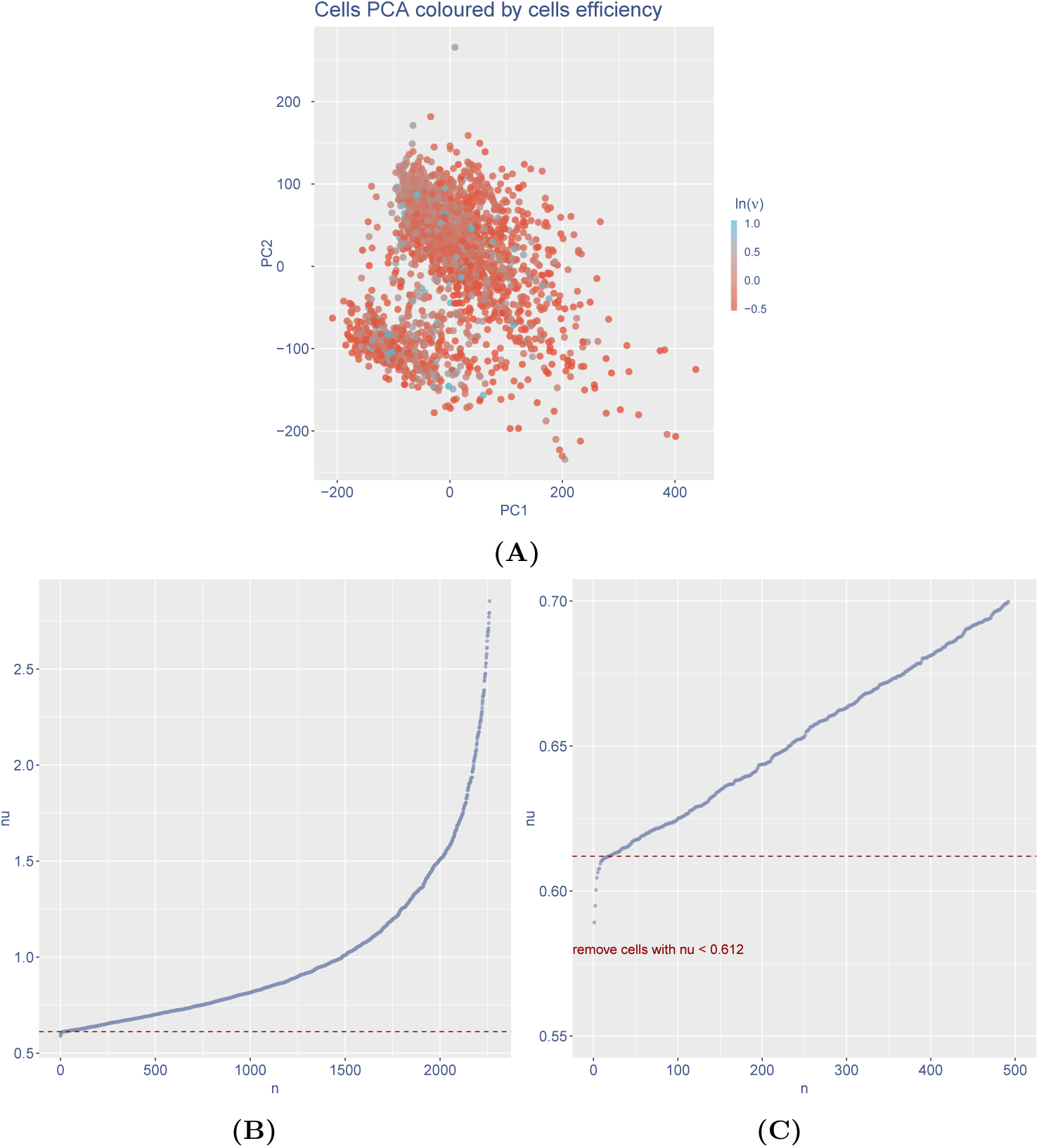
UDE plots for data cleaning. **A**. PCA plot of the cells colored by estimated UDE; in this case they show uniform distribution, so COTAN assumptions are reasonable. **B**. Plot of ordered UDE values for all cells. **C**. Magnification of the previous plot showing the elbow point below which cells should be removed.

We assigned randomly each cell *c* to a type *t*(*c*) ∈ {1, 2, …, *K*}, (where *K* was 1, 6 or 15, depending on the dataset), with probabilities proportional to the cluster sizes. Then we generated the potential expression 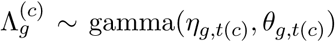 for each gene, with parameters depending on the gene and the cell type, and finally we generated the UMI counts 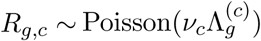.

The two homogeneous synthetic datasets contain a single type of cells, whose parameters were estimated from the largest cluster in the biological sample. Since in these datasets there is by construction no cell differentiation, but just variable cell extraction efficiency, all tests should result in accepting the null hypothesis, and hence they can be used to estimate the false positive rate and to check the distribution of the GPA’s *p*-value (see Figure 3C-D) and genome-wide GDI under the null hypothesis (see Figure 4C-D).

### 3.6 Gene co-expression space

GPA can be used to represent genes in a low-dimensional space and cluster them by co-expression. We already pointed out that two genes are shown to be co-expressed when their COEX is large, but if the aim is to cluster genes according to their differential expression, this information is not rich enough to infer something precise about their distance. For example it would be nice to include the information that both genes have strong negative COEX against a third gene. To be able to build a proper distance, we generalize this idea by comparing the whole co-expression patterns of the two genes against other genes: if these turn out to be similar, then the two genes should be probably considered “close” in the co-expression space.

The procedure starts by fixing a set *V* of genes to use as variables and a set *U* of genes to use as units to analyze (Figure 6). The two sets may be equal, but it is not required. Both sets, but in particular *V*, should contain only genes that are known or suspected to be differentially expressed in the sample, to reduce noise and improve quality.

To measure co-expression we consider the GPA COEX matrix for these genes. Applying the transform tanh 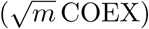 yields roughly binary values, since for many components 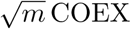 will have large absolute value^3^. There will be components near *±*1 for statistically significant joint and disjoint expression, and components randomly spread in (−1, 1) for the other couples. In this way the Euclidean distance between points becomes similar to a Manhattan distance for the co-expression pattern and hence these transformed matrix can be used as input for dimensionality reduction techniques and clustering.

The space in which genes *g* are represented with components *g* = (*g*_*v*_)_*v*∈*V*_

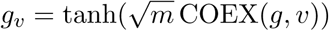

for all genes *v* ∈ *V* is the *co-expression space* relative to the set *V*. This space is suitable to perform dimensionality reductions, graphical representations and cluster analysis, as its Euclidean geometry is based on the gene coexpression patterns with genes in *V*. The choice of *V* will affect all these analysis and in fact a focus on peculiar subsets *V* may uncover information that is hidden or confounded in the genome-wide co-expression space (see also Section 2.3 of Results and Section 2.4 on the related concept of localized GDI).

### 3.7 COTAN pipeline

In the following subsections we describe the detailed COTAN pipeline (Figure 14). We developed and tested our method on drop-based techniques and in particular on 10X datasets and Drop-seq datasets. Analysis requires and starts from a matrix of raw UMI counts after removing possible cell doublets or multiplets and low quality or dying cells (with too high mtRNA percentage).

#### 3.7.1 Data cleaning

The first step is removing genes that are not significantly expressed (default threshold is 0.3% of cells) or unwanted (such as mitochondrial ones) to reduce the matrix memory size and to increase the analysis speed without losing information.

The user chooses either the linear or the square-root estimation framework (see Section 3.1.1), then begins an iterative procedure to filter out outlier cells (such as blood cells in a brain cortex dataset). In each iteration the UDE is estimated for all cells and UMI counts are simply normalized dividing by its value. Cells are then represented on the PCA plane and clustered by Mahalanobis distance (two clusters, *A, B*, complete linkage clustering). The clustering algorithm will detect outlier cells which will fall into the smallest cluster *B* (Figure 15A). A subsequent plot displays the most abundant genes expressed in *B*, to allow the user to check if they are peculiar in any way (Figure 15C). The user may choose to drop the cells in *B* and do another iteration, or to stop the procedure, when the PCA plot does not show obvious outliers (Figure 15B-D).

After the last iteration there are two final quality checks on the estimated UDE of cells. Firstly the PCA plot colored by UDE should not show a clear separation of cells with high and low UDE (Figure 16A). In fact COTAN builds on the assumption that UDE is not gene-dependent (see Section 3.1.1) and if the PCA plot is polarized by UDE, this may be false. Secondly, the plot of sorted UDE values will detect if the efficiency drops notably for a small fraction of cells. If this is the case, we usually want to exclude cells below the elbow point (Figures 16B-C, notice that UDE values are normalized to have mean 1, so there is not absolute threshold for acceptable efficiency). If cells are removed, another estimation iteration is due.

#### 3.7.2 Co-expression tables

This is a genome-wide procedure to compute the tables of observed counts for each pair of genes (see Section 3.2). If *n* is the number of genes in the sample, four *n* × *n* matrices store the number of cells in each of the four conditions (expressing both genes, only the first one, only the second one or none). Constitutive genes that appear in every single cell cannot be used and are removed in this step (saving a list of them).

#### 3.7.3 Expectation of co-expression table values

Before computing the expected values for the co-expression tables, it is necessary to estimate the dispersion parameter *a*_*g*_ of each gene *g* following the mathematical framework of Section 3.1. The approximated values are determined iteratively by simple bisection. Genes for which the estimated dispersion would be negative are instead modelled with zero dispersion and mean expression increased to (1 + *b*_*g*_)*λ*_*g*_. The positive parameter *b*_*g*_ is encoded as −*a*_*g*_ so that one single parameter can account for both cases. The fraction of genes with negative *a*_*g*_ is reported.

Afterwards the tables of expected values are computed like the observed counts, as four *n* × *n* matrices corresponding to the same four conditions.

#### 3.7.4 GPA *p*-values and COEX matrix

The script computes, for each pair of genes, the GPA test statistics *S*, the corresponding *χ*^2^(1) *p*-value and COEX index (Section 3.3) and saves them in three *n* × *n* matrices. The last two files are the primary output of COTAN analysis.

### 3.8 Seurat pipeline for Figure 1

Seurat (3.1.0) workflow was performed on E16.5 hippocampal dataset^16^ following the Guided Clustering Tutorial^29^, with modifications. Data import (CreateSeuratObject) was done using min.cells 3 and min.features 200. The selected range for the number of features was between 200 and 4000; the maximum allowed fraction of mitochondrial genes per cell was 7.5%. Normalization was done using the default parameters. The correlation was then calculated on the whole Seurat normalized data matrix and the heatmap was plotted subsetting this (Figure 1B). Figure 1D was plotted by calling the function `FindVariableFeatures` with selection method VST.

## 4 Conclusion

In conclusion, we introduced a new method for the analysis of scRNA data and described its application to datasets of embryonic hippocampus and cerebral cortex, which show high and documented cell identity diversity. We found that COTAN is well suited to identify gene pairs which are jointly or disjointly expressed in single cell. Moreover, COTAN GDI is a new parameter to assess differentially expressed genes in a dataset.

Finally, COTAN could cluster genes with consistent co-expression in single cell rather than cells with consistent gene global expression. Notably, COTAN features makes it a novel tool that, unlike previous indirect approaches^30^, can directly infer single-cell gene interactome from scRNA-seq dataset.

## Footnotes

1 The biological expression is here considered random, with unknown distribution of mean *λ*_*g*_, and measured not in real molecules, but in expected reads, taking the extraction efficiency into account.

2 The test explained above can give statistical significance to such a hypothesis.

3 This is because its square is the statistics *S* from GPA test for independence (see Section 3.3).

